# The interactome of *Cryptococcus neoformans* Rmt5 reveals multiple regulatory points in fungal cell biology and pathogenesis

**DOI:** 10.1101/2022.01.13.475903

**Authors:** Murat C. Kalem, Harini Subbiah, Shichen Shen, Runpu Chen, Luke Terry, Yijun Sun, Jun Qu, John C. Panepinto

**Author notes:** Corresponding author: John C. Panepinto, Department of Microbiology and Immunology, Witebsky Center for Microbial Pathogenesis and Immunology, Jacobs School of Medicine and Biomedical Sciences, 955 Main Street, Room 5229, University at Buffalo, SUNY, Buffalo, NY 14203, **Email:**.

## Abstract

Protein arginine methylation is a key post-translational modification in eukaryotes that modulates core cellular processes, including translation, morphology, transcription, and RNA fate. However, this has not been explored in *Cryptococcus neoformans*, a human-pathogenic basidiomycetous encapsulated fungus. We characterized the five protein arginine methyltransferases in *C. neoformans* and highlight Rmt5 as critical regulator of cryptococcal morphology and virulence. An *rmt5*Δ mutant was defective in thermotolerance, had a remodeled cell wall, and exhibited enhanced growth in an elevated carbon dioxide atmosphere and in chemically induced hypoxia. We revealed that Rmt5 interacts with post-transcriptional gene regulators, such as RNA-binding proteins and translation factors. Further investigation of the *rmt5*Δ mutant showed that Rmt5 is critical for the homeostasis of eIF2α and its phosphorylation state following 3-amino-1,2,4-triazole-induced ribosome stalling. RNA sequencing of one *rmt5*Δ clone revealed stable chromosome 9 aneuploidy that was ameliorated by complementation but did not impact the *rmt5*Δ phenotype. As a result of these diverse interactions and functions, loss of *RMT5* enhanced phagocytosis by murine macrophages and attenuated disease progression in mice. Taken together, our findings link arginine methylation to critical cryptococcal cellular processes that impact pathogenesis, including post-transcriptional gene regulation by RNA-binding proteins.

**Significance:** The fungal pathogen *Cryptococcus neoformans* is a huge threat for people living with immune deficits, especially HIV/AIDS. Its virulence potential is dependent on virulence factors, stress adaptation, and thermotolerance. Post-transcriptional gene regulation is important for these pathogenic processes, but the mechanisms that govern post-transcriptional regulator function are unexplored. Protein arginine methylation is a major modification of post-transcriptional regulators that has not been investigated in pathogenic fungi. Here we investigated the role of arginine methylation by arginine methyltransferases on the biology and virulence of *C. neoformans*. Phenotypic characterization of deletion mutants revealed pleiotropic functions for RMTs in this pathogen. Further investigation of the Rmt5 interactome using proximity-dependent biotinylation revealed interactions with RNA binding proteins and translation factors, thereby impacting virulence-associated processes.

## Introduction

All eukaryotes employ sophisticated molecular mechanisms that modulate signaling and gene regulatory processes. Fine-tuning of these complex molecular processes enables organisms to respond to dynamic environmental changes. For example, post-translational modifications are fast and reversible and can drastically alter proteins either by activating or suppressing function or by catalyzing transitions between functional states (1–5). One type of post-translational modification is the methylation of arginines on histones and non-histone targets such as RNA-binding proteins (RBPs), which is catalyzed by arginine methyltransferases (RMTs) (6–12).

RMTs are evolutionarily conserved among eukaryotes, with diverse functions across kingdoms. Dysregulation of RMTs in humans is often associated with devastating disease pathologies such as cancer and neurodegeneration (13–15). Humans have nine RMTs with unique functions in a plethora of cellular processes, including transcription, splicing, and translation (16–19). For example, type 1 RMTs (Rmt1, -2, -3, -4, -6, and -8) catalyze asymmetric dimethylarginine, and type 2 RMTs (Rmt5 and Rmt9) catalyze symmetric dimethylarginine. Human Rmt7 is the only type 3 enzyme and solely catalyzes monomethylarginine. *Saccharomyces cerevisiae* has three RMTs — Hmt1 (human Rmt1 ortholog), Hsl7 (human Rmt5 ortholog), and Rmt2. Hsl7, is important for normal budding and cell morphology by interacting with the septin ring components. Hsl7 also methylates histones, thereby repressing transcription (20, 21). Hmt1 binds to genomic regions near tRNAs and snoRNAs (22) to regulate the transcription of noncoding RNAs, possibly through histone modifications and methylation of proteins involved in RNA polymerase complexes. Yeast Hmt1 and fission yeast Rmt3 methylate the ribosomal protein Rps2 and modulate ribosome biosynthesis (23, 24).

Fungi have a comparatively limited diversity of RMTs: *Candida albicans, Aspergillus nidulans*, and *Neurospora crassa* each harbor three. *Candida albicans* RMTs regulate the nuclear export of nucleocytoplasmic RBP Npl3 (25), whereas the *Aspergillus nidulans* RmtC (an Rmt5 ortholog) is crucial for thermotolerance and resistance to hydrogen peroxide. In the plant pathogenic fungi *Magnaporthe oryzae* and *Penicillium expansum*, arginine methylation regulates their development and pathogenicity (26, 27). The potentially fatal human-pathogenic *Cryptococcus neoformans* harbors five RMTs, but very little is known about these. Our previous work established the importance of post-transcriptional and translational regulation in the virulence and pathogenicity of *C. neoformans* as well as antifungal resistance (28–33). Here, we characterized the functional roles of its RMTs to investigate how they fine-tune those regulatory mechanisms and influence the pathogenicity of *C. neoformans*. We analyzed stress responses, cell wall composition, and virulence of the fungus and identified Rmt5 as a key regulator of traits important for virulence. We then used proximity-mediated biotinylation protein interaction network analysis and RNA sequencing to further characterize the role of Rmt5 in modulating fungal cell biology. These findings are significant because RMTs are pharmacologically targetable and could expand the antifungal therapeutic toolbox.

## Results

### RMTs are evolutionarily conserved in fungi

We investigated the complement of RMTs in fungi by utilizing BLAST with the nine human methyltransferase sequences as queries and found five homologs in *C. neoformans* (Fig. 1a). Among the fungal species investigated, *C. neoformans* has the most RMTs according to conservation analyses. The presence of methyltransferase domains and conserved catalytic residues increased our confidence that these methyltransferases in *C. neoformans* have conserved catalytic functions (Fig. 1b). The homology of the five RMTs in *C. neoformans* indicates they can catalyze monomethyl and RMTs 1-4 can catalyze asymmetric dimethyl arginine methylation. Only Rmt5 is predicted to catalyze symmetric dimethylarginine methylation. We thus investigated further the phylogenetic conservation of Rmt5 across fungi from different phyla (Fig. 1c). To our knowledge, all fungal species harbor an Rmt5 ortholog, indicating that this enzyme is important and fundamental to fungal biology. Additionally, the Rmt5 catalytic domain, highlighted in yellow in Fig. 1c, is conserved across all investigated species. Nevertheless, experimental evidence is lacking to conclude that these proteins have conserved regulatory targets in fungal biology and similar roles in pathogenicity.

**Figure 1.**
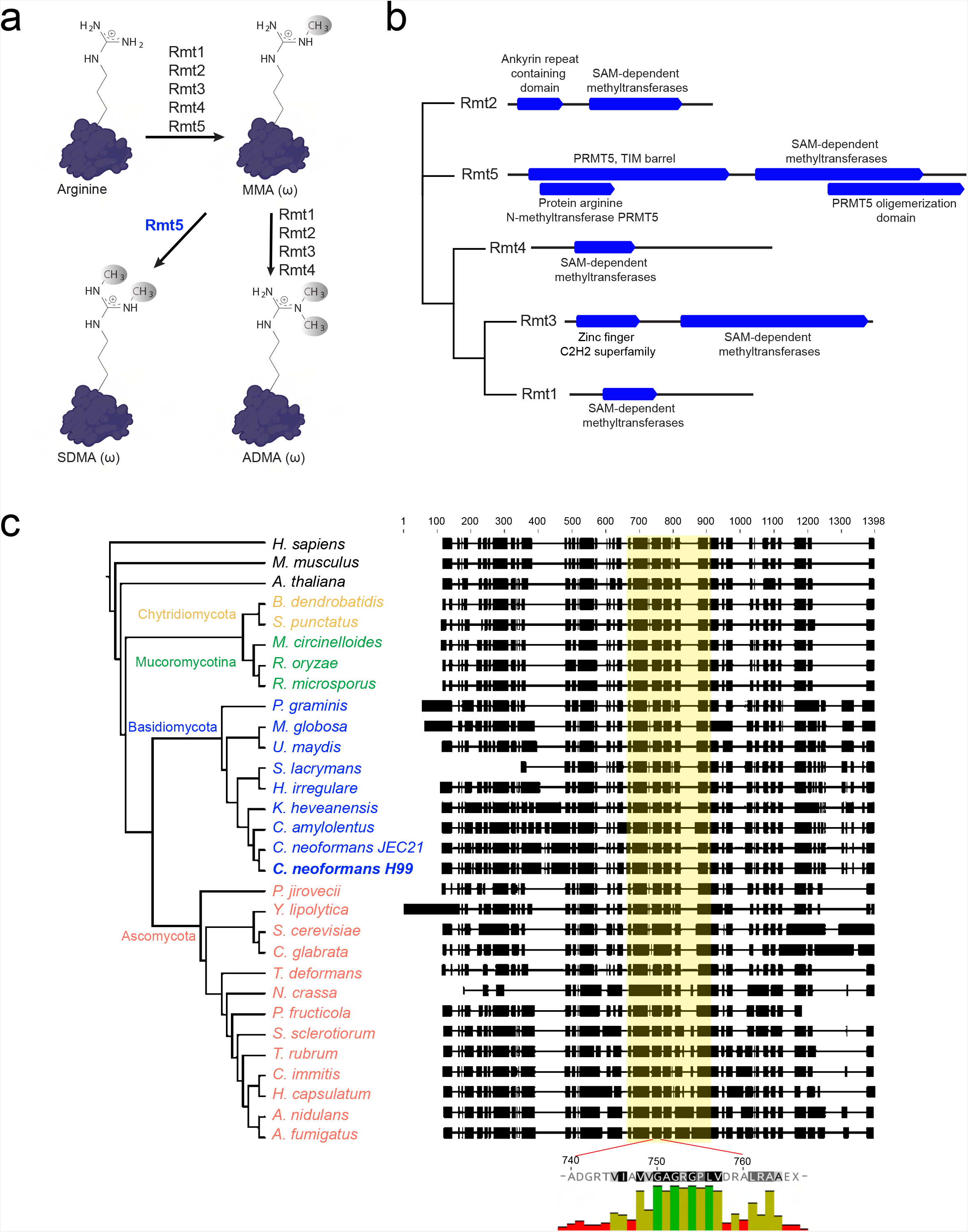
Phylogenetic classification of *C. neoformans* RMTs. (**a**) Schematic representation of *C. neoformans* RMTs and the methyl products they catalyze predicted by homology to human and mouse RMT protein sequences. (**b**) Conserved domains of *C. neoformans* RMTs as annotated by InterPro Domain search, which indicated the presence of catalytic and functional domains. (**c**) Phylogenetic comparison of Rmt5 across fungi. Rmt5 is evolutionarily conserved across fungi from different phyla with a highly conserved catalytic domain.

### Rmt5 regulates aspects of fungal cell biology involved in virulence

To investigate the roles of *C. neoformans* RMTs, we generated gene deletion mutants for each of the five homologs. None of the deletions were lethal, indicating that the genes are not essential for viability. We performed a phenotypic screen using a spot dilution assay to broadly identify deletion mutant phenotypes that may regulate virulence. Initial analyses revealed that the *rmt5*Δ is important for thermotolerance at 37°C and 38°C (Fig. 2a), an important virulence factor for many pathogenic fungi. We then investigated cell morphology at 30°C and 37°C, which revealed that the defect in thermotolerance stems from improper budding and failure of bud separation (Fig. 2b).

**Figure 2.**
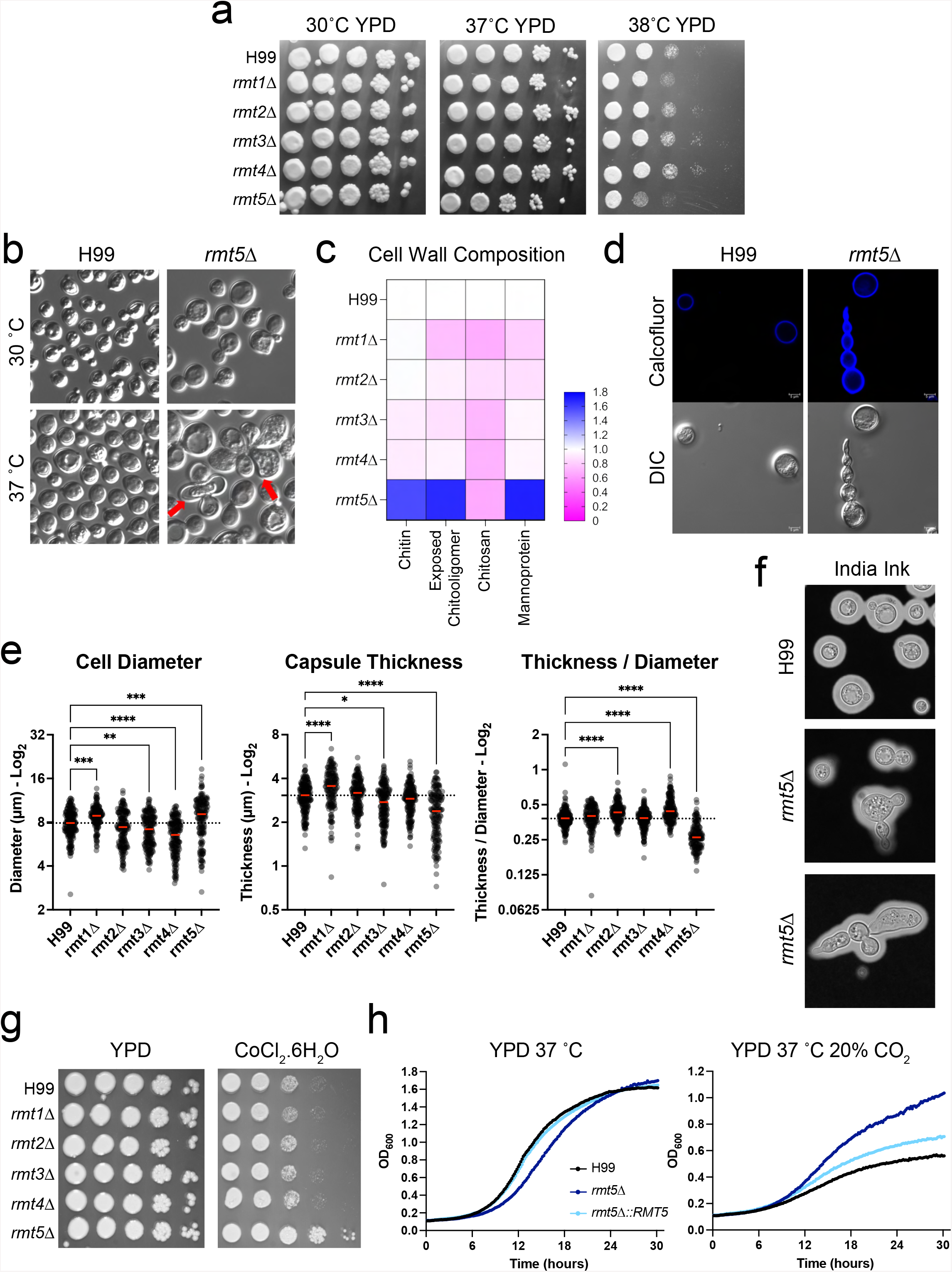
Functional characterization of RMTs shows that Rmt5 regulates various virulence-related features of fungal cell biology. (**a**) Spot plate analysis at different temperatures; plates were photographed after 72 h of incubation under the indicated conditions. Images are representative of three independent experiments. (**b**) Differential interference contrast images to show the morphology of cells grown overnight at the indicated temperatures. (**c**) Cells were grown overnight at 37°C, and different cell wall components were stained using calcofluor (chitin), wheat germ agglutinin (exposed chitooligomer), eosin Y (chitosan), and concanavalin A (mannoproteins). Abundances of cell wall components were quantified by flow cytometry; heat map represents the mean fluorescence intensity of each stain from 3 biological replicates relative to the wild type (H99). (**d**) Hemolymph was collected from *G. mellonella* larvae infected with either the H99 or *rmt5*Δ strain to asses cell morphology and chitin content via calcofluor staining. (**e**) Cells were grown under capsule-inducing conditions and stained with India ink to measure capsule thickness and cell diameter. Red bar represents the mean from 3 biological replicates. \**p* < 0.05; \*\**p* < 0.01, \*\*\**p* < 0.001, **** *p* < 0.0001 using a one-way ANOVA and Tukey’s multiple comparisons test. (**f**) Representative India ink images showing a minor defect in capsule thickness along with the signature morphological defect of the *rmt5*Δ cells. (**g**) Spot plate analysis under CoCl_2_-nduced hypoxia. (**h**) Growth under 20% CO_2_.

The cryptococcal cell wall and polysaccharide capsule are two critical elements of cell biology that are also involved in virulence. We investigated the cell wall composition of strains grown in yeast extract-peptone-dextrose (YPD) medium at 37°C. The *rmt5*Δ mutant had a remodeled cell wall containing increased levels of chitin, exposed chitooligomers, and mannoproteins. Deletions of the other RMTs resulted in minor decreases of these cell wall components (Fig. 2c). We also analyzed the cell wall composition of fungal cells isolated from the hemolymph of infected *Galleria mellonella* larvae. Calcofluor staining confirmed that the increase in chitin content is recapitulated in an *in vivo* infection model (Fig. 2d). There were also minor changes in capsule thickness under capsule-inducing conditions: the *rmt5*Δ mutant exhibited a modest but significant decrease in the capsule thickness-to-cell diameter ratio, whereas *rmt2*Δ and *rmt4*Δ mutants exhibited increases (Fig. 2e, f).

Adaptation to hypoxic environments and CO_2_ are important for environmental fungi that infect humans, and hypoxia responses are critical for *C. neoformans* virulence (34, 35). Therefore, we performed a spot dilution assay in the presence of cobalt chloride (CoCl_2_) and discovered that the *rmt5*Δ mutant is resistant to chemically induced hypoxia (Fig. 2g). We then investigated the growth rate in the presence and absence of 20% CO_2_. Results revealed that the *rmt5*Δ mutant was also resistant to high CO_2_ levels (20%) and grew better than the wild type under this condition (Fig. 2h). Overall, these findings indicate that Rmt5 regulates aspects of fungal cell biology that are important for virulence, with Rmt5 deficiency both promoting (resistance to hypoxia) and attenuating (thermosensitivity, reduced capsule thickness, and cell wall defects) pathogenic properties.

### Rmt5 interactome reveals effectors in post-transcriptional and translational regulation

Using a green fluorescent protein (GFP)-Rmt5 fusion protein, we observed that Rmt5 accumulates in the bud neck and future bud sites of *C. neoformans* cells (Fig. 3a), consistent with the *rmt5*Δ-induced budding defect. To characterize the molecular pathways that Rmt5 is involved in, we took an open-ended approach with proximity-dependent biotin labeling. We tagged Rmt5 with a C-terminal TurboID-3×Myc tag and expressed this construct in the *rmt5*Δ mutant cells (Fig. 3b). When grown at 37°C to mid-log growth stage, 299 biotinylated proteins were identified in two of three biological replicates, with 10% protein coverage and at least 2 spectral counts (see Table S2).

**Figure 3.**
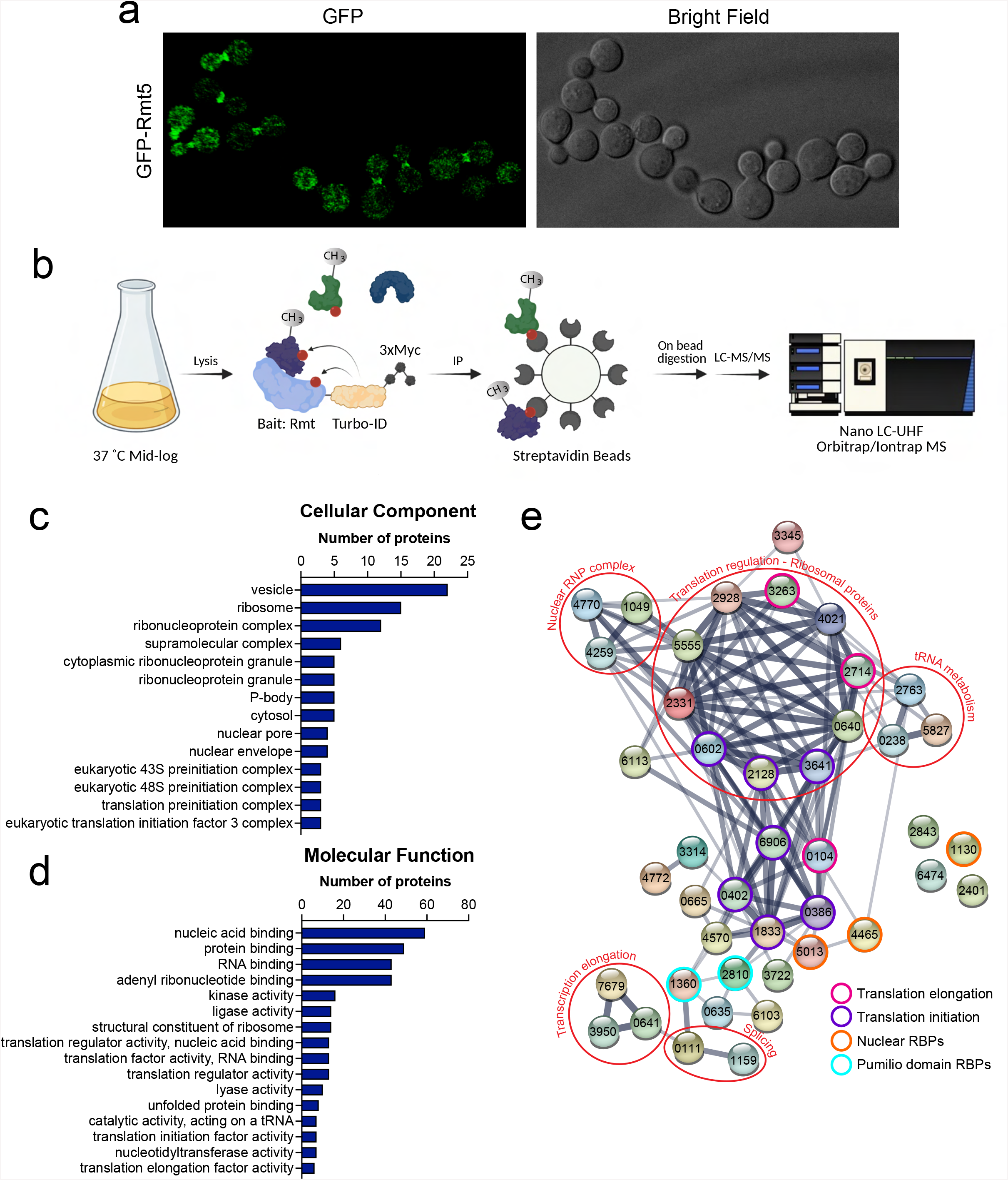
Rmt5 interactome includes effectors involved in post-transcriptional gene regulation. (**a**) Images of cells expressing Rmt5 tagged with GFP and grown at 37°C to mid-log stage. (**b**) Diagram summarizing the experimental design for the proximity-dependent biotinylation to determine the Rmt5 interactome. Biotinylated proteins were identified from 3 biological replicates using LC-MS/MS. (**c, d**) Gene ontology analysis of the interactome proteins; 299 interactome proteins were analyzed for enriched gene ontology terms on FungiDB. Graphs show the numbers of proteins in each cellular component and molecular function ontology category. Only categories that are enriched with a *p* value of 0.05 or better were selected. (**e**) RNA-binding protein functional association network. Proteins identified in the “RNA binding” category in the molecular function ontology search were investigated using String-DB. FungiDB gene identifiers (excluding CNAG_0) are presented within each node.

We analyzed the 299 interacting proteins for enriched gene ontology categories using FungiDB. We focused on cellular components and molecular functions with a *p* value threshold of 0.05. Gene ontology analysis revealed that Rmt5 interactome proteins are present in many cellular components, including ribosomes, ribonucleoprotein complexes, processing bodies, the nuclear pore, and nuclear envelope (Fig. 3c, Table S3). Consistent with this, enriched molecular function gene ontology terms included nucleic acid binding, RNA binding, kinase activity, and functions involved in translation regulation (Fig. 3d). Because RBPs such as human Rmt5 are key effectors of post-transcriptional gene regulation, we generated a protein interaction network of RBPs from the RNA-binding molecular function gene ontology category (Fig. 4e), which included translation initiation and elongation factors and RBPs with pumilio, La, and musashi domains as well as factors related to transcription elongation, splicing, and the nuclear envelope. Overall, our Rmt5 interactome analysis, while pioneering the use of TurboID in pathogenic fungi, identifies a role for Rmt5 in multiple molecular steps central to eukaryote biology.

**Figure 4.**
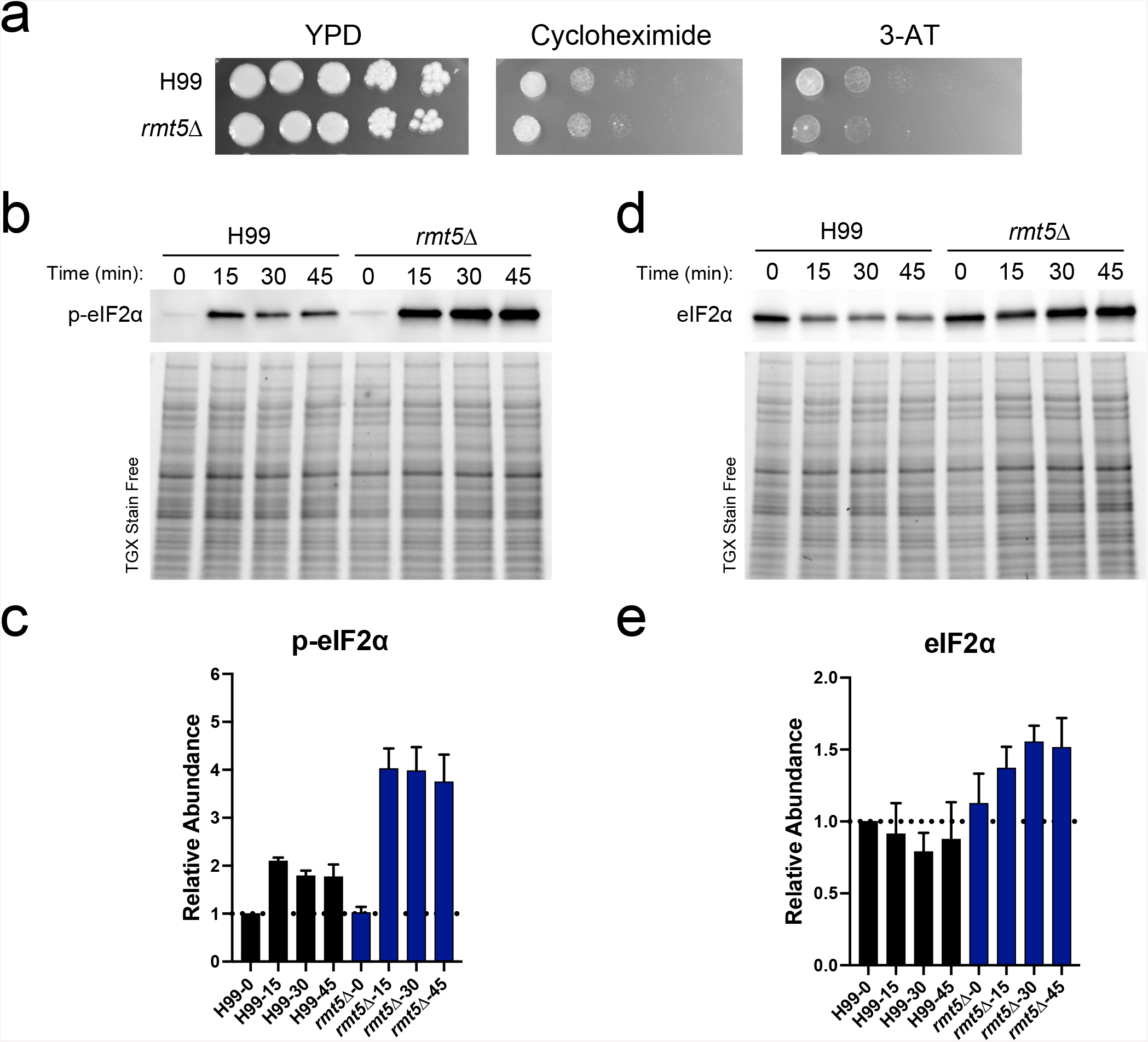
Rmt5 modulates translation by regulating eIF2α abundance and phosphorylation during 3-AT-induced ribosome stalling. (**a**) Spot plate analysis using translation inhibitors. Cells were grown on agar plates containing translation elongation inhibitors cycloheximide (250 ng/mL) and 3-AT (40 mM); photographs were taken after 3 days of incubation and are representative of three biological replicates. (**b**) Immunoblot showing the p-eIF2α abundance in a 3-AT time course. Cells were collected every 15 min following 3-AT treatment. p-eIF2α levels are normalized to total protein at the indicated time points. (**c**) Normalized quantification of p-eIF2α relative to H99 at 0 min (H99-0). Immunoblot (**d**) and quantification (**e**) of the abundance of total eIF2α. Error bars represent the SEMs from 3 biological replicates.

### Translation regulation in the *rmt5*Δ mutant is driven by altered eIF2α homeostasis

Because the Rmt5 interactome included many translation initiation and elongation factors, we investigated the response of the *rmt5*Δ mutant to translation inhibitors. Although the mutant exhibited wild-type growth in a spot dilution assay in the presence of cycloheximide, which inhibits eEF2-mediated tRNA translocation (36), the *rmt5*Δ mutant was sensitive to 3-amino-1,2,4-triazole (3-AT), which stalls ribosomes at histidine codons (37, 38) (Fig. 4a).

3-AT-induced histidine starvation and consequent ribosome collisions repress translation initiation by promoting the phosphorylation of eukaryotic translation initiation factor 2 (eIF2α) via kinase Gcn2 (39). Thus, we measured the levels of eIF2α and phosphorylated (p)-eIF2α during 3-AT treatment. Western blotting revealed that the treatment increased p-eIF2α levels 2-fold in wild-type cells but increased levels 4-fold in *rmt5*Δ cells (Fig. 4b, c). Interestingly, the levels of total eIF2α under the same conditions were higher in *rmt5*Δ cells than in the wild-type controls (Fig. 4d, e). This increase of total eIF2α levels alone cannot account for the increase in the p-eIF2α levels, suggesting that Gcn2 kinase activity was increased in *rmt5*Δ mutants. Overall, these findings indicate that Rmt5 is involved in the homeostasis of eIF2α and its phosphorylation.

### Rmt5-mediated transcriptome remodeling

Effectors of post-transcriptional gene regulation, such as RBPs, were among the most represented groups of proteins in the Rmt5 interactome. We reasoned that the transcriptome would be altered in cells lacking the Rmt5-mediated regulatory modifications. We employed RNA sequencing to investigate the global changes to the transcriptome in cells grown at 37°C to mid-log growth stage. We identified 310 genes that were upregulated and 69 that were downregulated in the *rmt5*Δ mutant, with adjusted *p* values of 0.05 and a 1.75-fold change cutoff (Fig. 5a, Table S4). Genome-wide mapping of all upregulated transcripts revealed that the majority (201 of 310 transcripts) belonged to chromosome 9 (Fig. 5b), which comprises 580 genes. This suggested to us that the clone of *rmt5*Δ used in the RNA-seq experiment harbored a chromosome 9 aneuploidy. To confirm this, we performed quantitative PCR using genomic DNA from wild-type and *rmt5*Δ cells, using the abundance of *CDC24* to assess chromosome 9 dosage. Our results confirmed a chromosome 9 duplication in the *rmt5*Δ clone used (Fig. 5c). However, the duplication was not present in the complemented strain.

**Figure 5.**
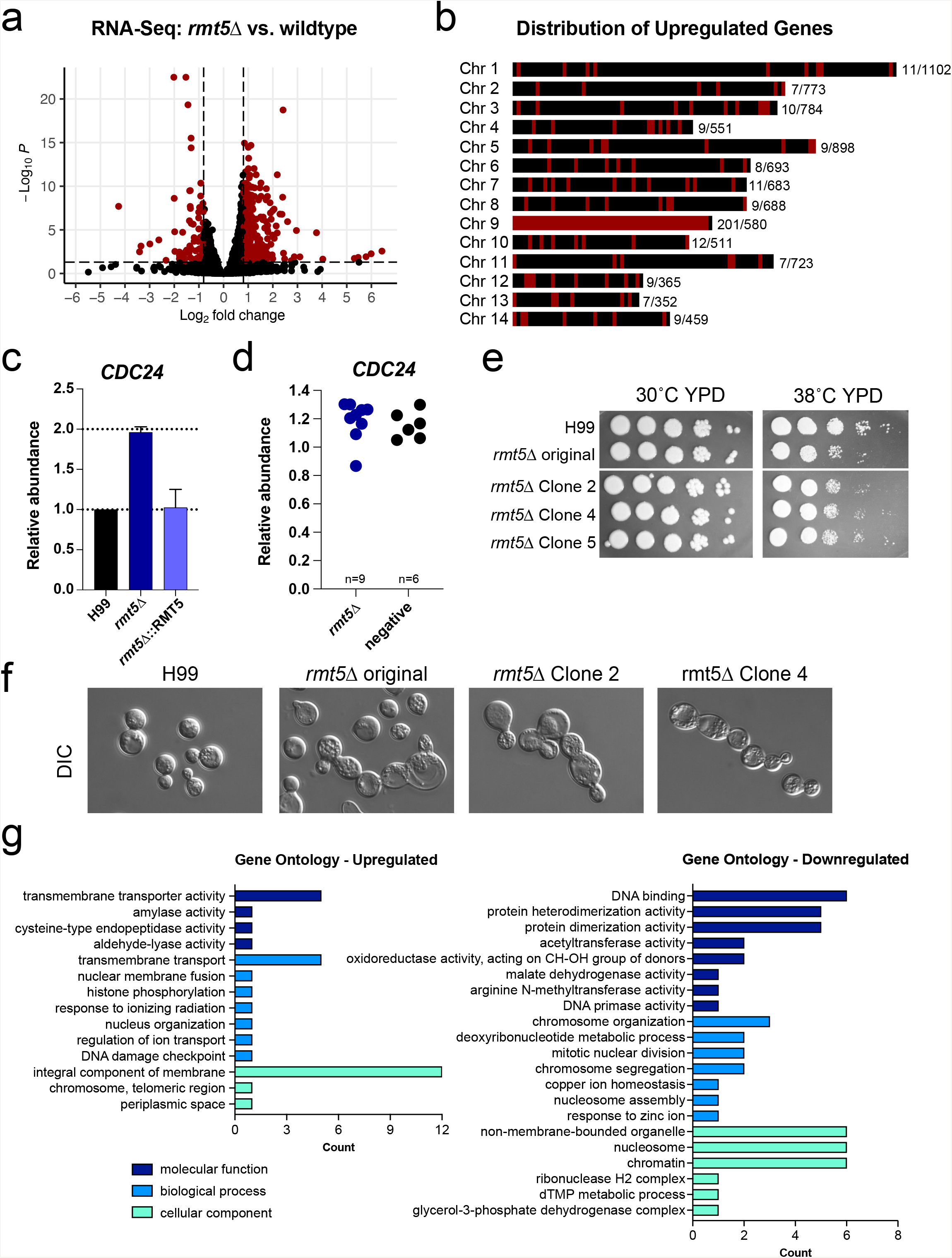
Rmt5-mediated transcriptome remodeling. (**a**) RNA-seq analysis of the transcriptome changes in the *rmt5*Δ. Volcano plot showing 69 downregulated and 310 upregulated transcripts in the *rmt5*Δ mutant with a 1.75-fold change and an adjusted *p* value threshold of 0.05. (**b**) Chromosome maps showing the distribution of all upregulated genes in the *rmt5*Δ mutant. Upregulated transcripts disproportionately belong to chromosome 9. (**c**) Genomic DNA from H99, *rmt5*Δ, and complement strains were used as templates for quantitative PCR reactions to investigate the abundance of chromosome 9 gene *CDC24*. Abundance of *CDC24* was normalized to *GRP7*, which is on chromosome 8. Error bars represent the SEM from 3 biological replicates. (**d**) Chromosome 9 dosage in nine new *rmt5*Δ mutant clones and six negative clones. Each dot represents an independent clone. (**e**) Spot plate analysis at 30°C and 38°°C confirming that the thermotolerance defect is not dependent on chromosome 9 aneuploidy. (**f**) The budding defect and morphological abnormalities observed in the *rmt5*Δ mutant at 37°C are not attributable to chromosome 9 aneuploidy. (**g**) Aneuploidy-independent transcriptome remodeling reveals enriched gene ontology categories related to chromatin, nucleosome, and cell membrane. Upregulated genes were filtered to exclude chromosome 9 genes and genes that encode ncRNAs. The remaining 45 genes were used as the input for the gene ontology term analysis. Downregulated genes were filtered to exclude ncRNAs, and the remaining 57 were used as the input for the gene ontology term analysis. Terms with a *p* value lower than 0.05 were selected. The full list of gene ontology terms is included in Table S5.

To further dissect the functional role of this chromosome 9 duplication and validate that the observed *rmt5*Δ phenotypes were attributable to the gene deletion, we generated nine additional independent *rmt5*Δ clones and analyzed chromosome 9 dosage; none exhibited the chromosome 9 aneuploidy observed in the original clone (Fig. 5d). We next compared the phenotype of the *rmt5*Δ clone harboring the aneuploid chromosome 9 with those of the euploid *rmt5*Δ clones and found that all exhibited similar defects in thermotolerance, cell morphology, and budding, confirming that these traits resulted from the loss of *RMT5* and were independent of the chromosome 9 aneuploidy (Fig. 5e, f). We also repeated the eIF2α immunoblots with two additional *rmt5*Δ clones, which confirmed that eIF2α dysregulation in the *rmt5*Δ mutant is likewise independent of chromosome 9 aneuploidy (Fig. S3)

We then excluded upregulated chromosome 9 genes from the RNA-seq data from the original clone, leaving 109 genes that were upregulated and 69 that were downregulated as a result of Rmt5-mediated transcriptome remodeling. Of note, 64 of the upregulated genes and 12 of the downregulated transcripts are classified as noncoding RNAs (ncRNAs), suggesting that Rmt5 regulates ncRNA expression. Gene ontology analysis of the protein-coding genes that were upregulated and downregulated yielded a small number of genes in each statistically enriched (*p* value threshold of 0.05) gene ontology category. Twelve of the upregulated genes are classified as integral part of cell membrane, which includes transporters, and the downregulated genes are in categories involved in DNA binding, chromosome organization, chromosome segregation, and nucleosome assembly (Fig. 5g, Table S5). We speculate that the downregulation of these genes contributed to the budding and morphology defects. Transcripts for histones H2A, H2B, H3, H4, and H1/5 were downregulated in the *rmt5*Δ mutant (Fig. S2). However, the abundances of H3 and H4 proteins did not differ significantly between the wild type and the *rmt5*Δ mutant (Fig. S2), suggesting that the impact of Rmt5 on histone mRNA levels does not affect the overall abundance of these proteins. The lack of large gene expression changes in the absence of *RMT5* suggests that Rmt5-mediated RBP regulation alters mRNA fate by regulating RNA processes that may or may not impact transcript abundance, such as RNA localization, RNP assembly, and translation.

### Rmt5 is important for cryptococcal pathogenicity

Lastly, we explored the role of RMTs in fungal pathogenesis using a *Galleria mellonella* model of infection (Fig. 6a). We found that larvae infected with the *rmt1*Δ fungal mutant exhibited attenuated disease progression. Surprisingly, even though the *rmt5*Δ mutant had various defects in phenotypes critical for virulence, we did not observe a virulence defect in the model. We then investigated the phagocytosis and killing of mutants by murine macrophages (Fig. 6b). The *rmt2*Δ, *rmt4*Δ, and *rmt5*Δ mutants were more readily phagocytosed, indicated by the higher percentage of infected macrophages. Killing analysis showed that the survival of *rmt1*Δ and *rmt5*Δ mutants was decreased after phagocytosis by macrophages. Because the *rmt5*Δ mutant was more likely to be phagocytosed and killed by murine macrophages, we reasoned that this mutant would have a virulence defect in mice. Therefore, we inoculated CBA/j mice intranasally with H99 and *rmt5*Δ mutant cells. The survival analysis revealed that the disease progression in mice infected with the *rmt5*Δ mutant was attenuated (Fig. 6c). This effect was attributed to the loss of *RMT5* because the probability of survival with infection by the complemented fungal strain was similar to that with the wild type. Lung sections from mice infected with the wild type or the mutant *C. neoformans* exhibited large regions of inflammation when stained with hematoxylin and eosin (Fig. 6d). Gomori methenamine silver staining of lung sections from mice infected with the *rmt5*Δ mutant revealed the budding defect of *C. neoformans* that was observed in culture (Fig. 6e). Overall, these findings indicate that RMTs, especially Rmt1 and Rmt5, are important for fungal pathogenesis and virulence.

**Figure 6.**
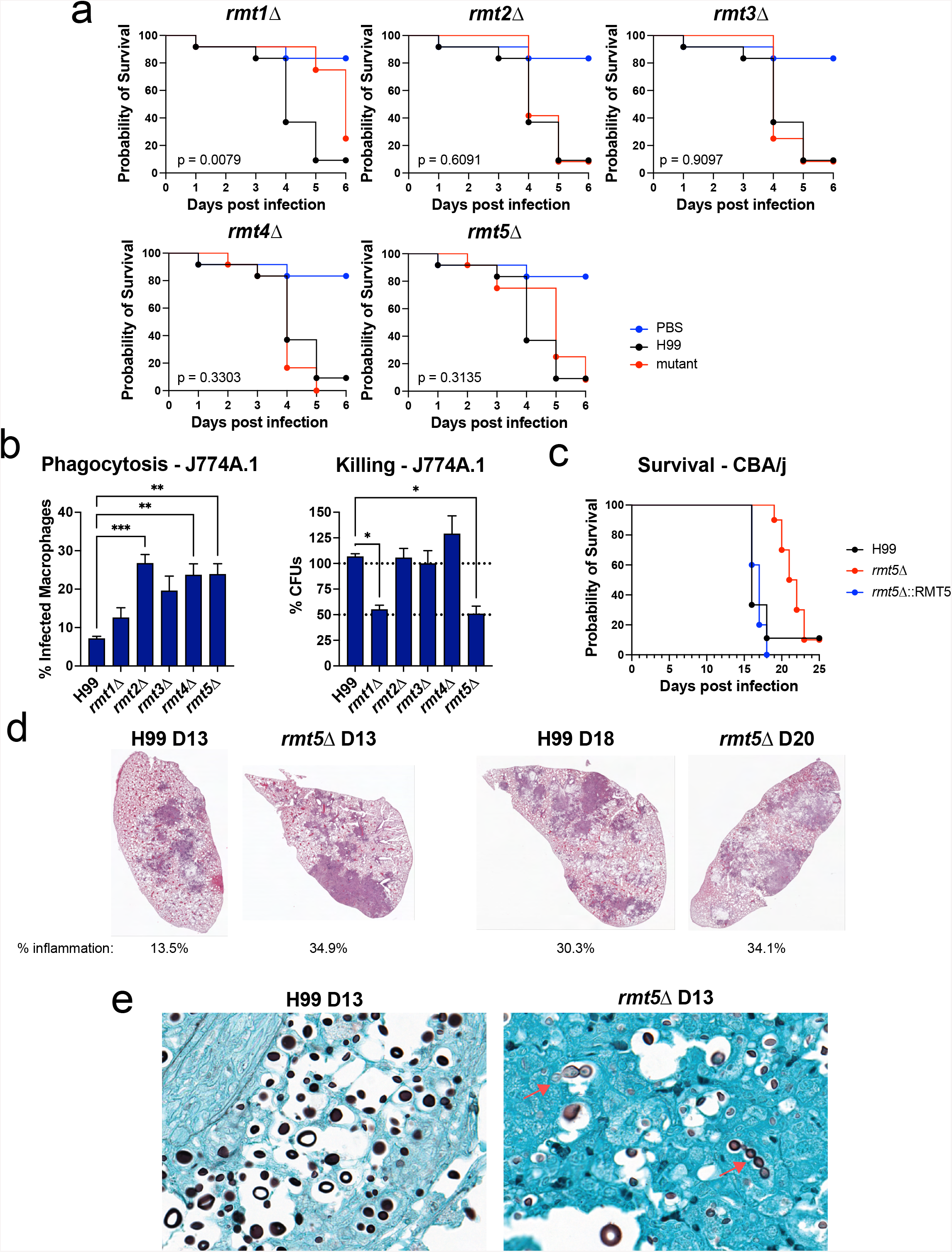
Role of RMTs in fungal pathogenesis. (**a**) RMT mutants were screened for pathogenesis defects using a *Galleria mellonella* larvae killing assay. Twelve larvae were infected per group; *p* values were calculated using a log rank (Mantel-Cox) test. The wild-type and PBS survival curves are duplicated on each graph for clarity; all infections were performed simultaneously. (**b**) Wild-type and mutant cells were co-incubated with activated J774A.1 murine macrophages. Phagocytosis of each strain was analyzed by flow cytometry. Killing was determined by analyzing the CFUs after phagocytosed fungi were co-incubated for 24 h with macrophages. Wild-type cells were assumed to survive 100%, and other strains are graphed relative to the wild type. Error bars represent SEMs. \**p* < 0.05; \*\**p* < 0.01, \*\*\**p* < 0.001 using a one-way ANOVA and Tukey’s multiple comparisons test. (**c**) H99, *rmt5*Δ, and *rmt5*Δ::*RMT5* strains were used to intranasally inoculate CBA/J mice (*n* = 10 per group for mutant strains, 9 for H99); *p* values were calculated using log rank (Mantel-Cox) test. (**d**) Hematoxylin and eosin staining of whole-lung sections. Inflamed areas, stained dark purple, were measured using QuantPath. Percent area with inflammation was calculated for each image. (**e**) Gomori methenamine silver staining highlighting the fungal cell morphology in brown. The *rmt5*Δ mutant-infected lung sections harbor yeast cells with the signature morphology and budding defect of the mutant.

## Discussion

*C. neoformans* is a human fungal pathogen that can cause fatal meningitis in patients with cell-mediated immunity defects, including transplant recipients and those living with HIV/AIDS, and is responsible for 15% of AIDS-related deaths (40–43). Treatment of *C. neoformans* infection is difficult because current treatments are not well tolerated and therapeutic options are limited, especially in resource-poor settings (44). Knowledge of the molecular and cellular processes that control fungal-specific biology and virulence-associated traits will serve as a foundation for future drug discovery efforts. Although post-transcriptional gene regulation in *C. neoformans* is known to impact virulence and drug resistance, the contribution of protein arginine methylation in this process has been a neglected area of study. However, the study of RMTs has the potential to reveal unique fungal molecular regulatory mechanisms that may aid the design of novel therapies, because these enzymes are pharmacologically targetable (45).

In this study, we began to characterize the functional roles of five *C. neoformans* RMTs by performing a broad phenotypic screen. Our screen identified Rmt5 as a regulator of key virulence-associated traits, including thermotolerance, capsule and cell wall regulation, and resistance to chemically induced hypoxia and high CO_2_. Notably, the *rmt5*Δ mutant exhibited temperature-dependent morphological and budding defects. Rmt5 was mainly localized to the cytoplasm but accumulated around the site of budding. To determine which proteins Rmt5 interacts with, we performed TurboID of biotinylated proteins, which can detect transient interactions, such as those between a methyltransferase and its substrate. Although the yeast Rmt5 ortholog, Hsl7, interacts with septin proteins, these were not identified as Rmt5 interactors in our TurboID screen. Instead, our screen identified fimbrin/Sac6, which also localizes to the bud neck of yeast and the hyphal tip collar in filamentous fungi. In addition, Cdc20, a known regulator of cytokinesis, was identified as an Rmt5 interactor. Further work is needed to define which of the Rmt5 targets are responsible for the thermosensitivity and the budding defect of the *rmt5*Δ mutant.

The Rmt5 interactome also included many RBPs and translation factors, suggesting a role for Rmt5 in post-transcriptional gene regulation. Such regulation can impact virulence-associated traits in *C. neoformans*. For example, deletion of Ccr4, the major cytoplasmic deadenylase, causes defects in thermotolerance and cell morphology that alter virulence potential (30, 46). One of the RBPs that interact with Rmt5 is the pumilio/FBF RBP family member Puf4. Our previous work showed that Puf4 is critical in post-transcriptionally regulating caspofungin resistance (28). Combined with our current Rmt5 interactome data set, we speculate that Rmt5 fine-tunes virulence- and antifungal resistance-associated cellular functions via methylating Puf4 and other effectors. Future work will investigate the role of arginine methylation of Puf4 in regulation of mRNA decay and antifungal resistance.

The methylation of ribosomal proteins is critical for the assembly of functional ribosomes in both human and yeast cells (47, 48). Arginine methylation is known to impact translation, and methylation of RGG motif-containing proteins represses translation by interacting with eIF4G in yeast. Scd6, an RGG domain protein, interacts with eIF4G and augments its translational repression. Scd6 methylation by Hmt1 (an Rmt1 ortholog) enhances this interaction, thereby regulating this repression. *C. neoformans* lacks an Scd6 homolog, but there likely are other RGG domain-containing proteins that replicate this function (49). Moreover, Rmt5 may regulate translation initiation via eIF4E or eIF4B. Indeed, the *rmt5*Δ cells exhibited hyperphosphorylation of eIF2α as well as a small increase in total eIF2α levels in the presence of 3-AT, which induces ribosome collisions that inhibit translation. This suggests that the absence of arginine methylation by Rmt5 results in dysregulation of eIF2α phosphorylation in response to pharmacologically induced stress. Alternatively, the absence of Rmt5-regulated translation may predispose ribosomes to collide in response to 3-AT. Nevertheless, our *C. neoformans* Rmt5 interactome included many translation initiation and elongation factors, implicating Rmt5 in translation regulation. We also know that translatome remodeling in *C. neoformans* is crucial for tolerance to temperature and compound stress (29, 30, 33, 50). Our work demonstrates that Rmt5 modulates translation under stress by targeting translation initiation and elongation factors, ribosomal proteins, and RBPs. The specific molecular mechanisms by which Rmt5 impacts the ribosome, its function, and its regulation are under investigation.

Rmt5 is a post-transcriptional regulator as defined by our study and others (51) and thus was expected to widely impact gene expression. Serendipitously, our RNA-seq analysis showed that many genes located on chromosome 9 were upregulated. Further investigation revealed that the *rmt5*Δ mutant we were testing had chromosome 9 aneuploidy. *C. neoformans* has high genome plasticity, and aneuploidies occur commonly in response to stress and antifungal drugs (52, 53). There are clinical isolates that have disomic chromosomes (54), and *C. neoformans* aneuploidy enables resistance to fluconazole and flucytosine (55–59). We first hypothesized that this aneuploidy might remediate the loss of Rmt5 or else cause the observed *rmt5*Δ phenotypes. Thus, we repeated the biolistic transformations and isolated independent deletion clones. These clones did not have the chromosome 9 aneuploidy but still exhibited defects in thermotolerance, cell morphology, budding, and eIF2α homeostasis following 3-AT treatment. We therefore concluded that the aneuploidy was not consequent to *RMT5* deletion to counteract the fitness defects. However, the downregulation of genes for DNA-binding proteins, chromosome organization, and chromosome segregation suggests that *C. neoformans* is predisposed to develop aneuploids in the absence of *RMT5*. This rather serendipitous observation of aneuploidy was possible because of the advances in public databases such as FungiDB and the genome view function (60). The genomic locations of genes found to be upregulated or downregulated in RNA-seq analyses are not often examined, yet this can provide critical hints about chromosome abnormalities. Indeed, this prompted us to perform a more rigorous investigation of Rmt5 function.

Histones, as part of the DNA binding and nucleosome gene ontology categories, were downregulated in the *rmt5*Δ mutant. Follow up experiments revealed that histone protein abundances are unaltered in the mutant despite the significant downregulation. Histone transcripts in humans and most higher eukaryotes lack a canonical poly-A tail, and instead contain a 3’-terminal stem loop structure that is recognized by stem-loop binding protein, SLBP1 (61–63). SLBP1 binding to 3’ stem loop of histone facilitates translation of histone mRNAs, taking the place of the poly-A tail (64, 65). The *C. neoformans* genome does not encode an SLBP1 ortholog, suggesting that other RBPs regulate histone mRNAs. Intriguingly, the presence of histone transcripts in our RNA-seq data sets using a poly-A RNA isolation method suggests that *C. neoformans* histone transcripts are polyadenylated. Histone mRNAs in plants are also polyadenylated, suggesting that histone mRNA regulation has diverged across eukaryotic evolution. In light of our Rmt5 interactome data and RNA-seq data, we further speculate that an Rmt5-targeted RBP might function in histone pre-mRNA processing and translation.

The characterization of cryptococcal RMT mutants revealed that both *RMT1* and *RMT5* are critical for pathogenesis in host organisms. Both *rmt1*Δ and *rmt5*Δ mutants showed enhanced phagocytosis and killing by murine macrophages. We reasoned that the cell morphology defects and exposed chitin-rich cell wall of the *rmt5*Δ mutant were responsible for the enhanced phagocytosis *in vitro* and attenuated virulence *in vivo*. Intriguingly, the *rmt1*Δ mutant did not exhibit thermotolerance or cell fitness defects in our screen. Overall, our study pioneers the characterization of RMTs in *C. neoformans*, defines the regulatory network of Rmt5, and proposes the key cellular processes modulated through this post-translational modification-driven network. Future work will focus on characterization of specific Rmt5-driven methylation events and their functional consequences while exploring the contributions of Rmt1 to *C. neoformans* biology, drug resistance, and pathogenesis.

## Materials and Methods

### Strains and molecular cloning

Strains used in this study were derived from *Cryptococcus neoformans* var. *grubii* strain H99, a fully virulent strain (gifted by Peter Williamson, UIC, NIAID) derived from strain H99O (gifted by John Perfect, Duke University). All primers used in this study are listed in Table S1. Strains were created using biolistic transformation. Gene deletions were generated using homologous recombination of deletion constructs. Deletion constructs include upstream and downstream homology sequences on both sides of the NAT resistance cassette. *RMT5-*TurboID-3×Myc and GFP-*RMT5* constructs were built using NEBuilder linear assembly. The TurboID-3×Myc sequence was amplified from pFB1434, obtained from François Bachand (Addgene plasmid no. 126050) (66). The *RMT5-*TurboID-3×Myc construct includes the native *RMT5* promoter, coding sequence excluding the stop codon, TurboID-3xMyc sequence, and *RMT5* terminator. The GFP-*RMT5* construct includes native *RMT5* promoter (∼1 kb), GFP sequence (no stop), *RMT5* coding sequence, and native *RMT5* terminator. Both NEBuilder constructs were PCR amplified from NEBuilder reactions and ligated into pJET1.2/Blunt. pJET1.2-*GFP-RMT5* and pJET-*RMT5-* TurboID-3×Myc were co-transformed with PCR-amplified NEO resistance cassettes into the *rmt5*Δ background to generate *rmt5*Δ::GFP*-RMT5* and *rmt5*Δ::*RMT5*-TurboID-3×Myc, respectively.

### Phylogenetic analysis

*C. neoformans* orthologs of RMTs were identified using a BLAST algorithm. Identified hits were confirmed using a reciprocal BLAST search. Five RMTs were compared using MUSCLE alignment followed by a neighbor-joining tree protein alignment on Geneious. A BLAST search was then used to find the orthologs of Rmt5 in fungal species from *Ascomycota, Basidiomycota, Mucoromycotina*, and *Chytridiomycota* phyla by using human PRMT5 (UniProt ID: O14744) as the query. Human and mouse PRMT5 were used as outgroups. All Rmt5 ortholog sequences were aligned using MUSCLE, and a phylogenetic tree was built using Geneious as described above.

### Spot plate dilution assay

Cells were grown overnight at 30°C in YPD broth. Overnight cultures were washed with sterile distilled water, and the optical density at 600 nm was adjusted to 1 in water. Adjusted cultures were 1:10 serially diluted 5 times, and 5 µL of each dilution was spotted onto YPD agar plates containing the selected drugs. Agar plates were incubated 2–3 days at the temperatures indicated in the text and photographed.

### Cell wall composition analysis

Cells were grown overnight at 37°C in YPD broth and then washed with sterile distilled water. The cells were stained with calcofluor white, wheat germ agglutinin (fluorescein isothiocyanate conjugated), eosin Y, and concanavalin A (Alexa Fluor 488 conjugated) as described previously (28) to detect cell wall chitin, exposed chitooligomers, chitosan, and mannoproteins, respectively. Flow cytometry data were acquired using a BD LSRFortessa cell analyzer and analyzed using FlowJo v10.0 software.

### *Galleria mellonella* infection

*G. mellonella* larvae were weighed, and larvae weighing 250 ± 25 mg were selected. Inocula were prepared from *C. neoformans* cells grown overnight at 30°C in YPD broth and washed twice in PBS. Ten microliters of the inoculum (1×10^8^ cells/mL) was injected into the haemocoel of each larva through the proleg using a Hamilton syringe after the injection area was disinfected with an alcohol swab. Each experimental and control group contained 12 larvae. One group was injected with PBS to serve as a control to monitor survival following physical injury. The larvae were incubated at 37°C and monitored daily to record survival. Death was recorded when the larvae melanized and/or failed to respond to physical stimuli. Survival curves were plotted over time to compare the survival of larvae infected with wild-type versus mutant *C. neoformans*. Curves were statistically analyzed using a log rank Mantel-Cox test, and *p* values were reported for each comparison.

### Capsule induction and analysis

Overnight cultures were grown at 37°C and washed with sterile distilled water. Cell pellets were resuspended in 5 ml RPMI medium, and 10 µL of the cell suspension was added to 3 mL RPMI medium in each well of a 12-well plate. Cells were incubated for 24 h at 37°C with 5% CO_2_. Cells were pelleted, stained with India ink, and imaged using a differential interference contrast filter. Cell size and capsule thickness were measured using Fiji (ImageJ).

### Analysis of phagocytosis and killing by macrophages

J774A.1 murine macrophages (gifted by Jason Kay, University at Buffalo) were routinely cultured in RPMI medium supplemented with 10% fetal bovine at 37°C with 5% CO_2_. Phagocytosis and killing assays were adapted from those described by Wormley et al. (67). Macrophages (1×10^5^ per well) were seeded on 96-well plates and incubated for 18–24 h and then activated by incubating with 10 nM phorbol myristate acetate for 1 h. *C. neoformans* overnight cultures were pelleted, washed twice with PBS, and then resuspended in RPMI medium. Approximately 10^6^ fungal cells/mL were stained with 25 µg/mL calcofluor white for 15 min and then opsonized with 1 µg/mL monoclonal 18B7 antibody for 1 h. One hundred microliters of opsonized *C. neoformans* was added to each well of the activated macrophage cultures for a 1-h incubation. Cells were then washed four times with PBS and processed either for phagocytosis or killing analysis.

For phagocytosis, cells were fixed and stained with phalloidin-Alexa Fluor 488, and the percent macrophage population that had phagocytosed *C. neoformans* cells was quantified. For killing, 100 µL RPMI medium was added to each well, and phagocytosed *C. neoformans* were incubated for 24 h. The macrophages were lysed by adding 200 µL sterile water and incubated at room temperature for 5 min. Killing of *C. neoformans* were assessed using EddyJet 2 spiral plater to determine CFU following a 2-day incubation using a ProtoCOL 3 colony counter (Synbiosis). Survival of mutant strains was analyzed relative to that of H99 (wild type), which was set at 100%. Three biological replicates with 3–4 technical replicates each were analyzed for both phagocytosis and killing assays.

### Mouse infections

CBA/j mice were obtained from Jackson Laboratories. *C. neoformans* overnight cultures were washed twice with PBS and counted using a hemocytometer. CBA/j mice were briefly anesthetized with isoflurane and inoculated intranasally with 5×10^5^ fungal cells in 50 µL PBS. Ten mice per group were infected and monitored daily according to IACUC guidelines. Survival was analyzed using a log rank Mantel-Cox test. H99- and a *rmt5*Δ mutant-infected mice were kept separate.

Histological evaluations of lung and brain tissues were performed 13 days post infection. Briefly, tissues were harvested at the time of euthanasia and fixed in 10% buffered formalin for 24–36 h and then transferred to 70% ethanol. Tissues were sectioned and stained with hematoxylin and eosin and Gomori methenamine silver at the Roswell Park Histology Core Facility.

### Affinity purification of biotinylated proteins

*RMT5*-TurboID-3xMyc and H99 cells were grown to mid-log stage at 37°C in 2 L YPD medium that contained 50 µM biotin. Cells were pelleted and resuspended in 2 mL RIPA buffer (50 mM Tris-HCl [pH 7.5], 150 mM NaCl, 1.5 mM MgCl_2_, 1 mM EGTA, 0.1% SDS, 1% NP-40, 0.4% sodium deoxycholate, 1 mM dithiothreitol (DTT), 1 mM phenylmethylsulfonyl fluoride, 1× cOmplete, and 1× PhosSTOP). Cell suspensions were frozen in liquid nitrogen as droplets. Frozen pearls were then lysed using a coffee grinder (2 min) and with a mortar and pestle (2 min). The lysate was kept cold during this process by adding liquid nitrogen when necessary. Lysate powder was collected from the mortar, transferred to a conical tube, and resuspended in 10 mL RIPA buffer containing 0.4% SDS. Lysates were treated with 500 U Benzonase to digest RNA and DNA. Clarified lysates were then incubated with 100 µL streptavidin-Sepharose for 4 h at 4°C. Then, beads were washed four times with RIPA buffer containing 0.4% SDS and four times with 20 mM ammonium bicarbonate. Beads were resuspended in ammonium bicarbonate and stored at −80°C until liquid chromatography tandem mass spectrometry (LC-MS/MS) analysis.

### Protein digestion

A surfactant-aided precipitation/on-bead digestion protocol was employed according to a previous publication (68). Pelleted beads were resuspended with 100 μL 0.5% SDS, reduced by 10 mM DTT at 56°C for 30 min, and alkylated by 25 mM iodoacetamide at 37°C in darkness for 30 min. Both steps were performed with rigorous vortexing in a thermomixer (Eppendorf). Protein was precipitated by adding 6 volumes of chilled acetone with constant vortexing, and the mixture was incubated at −20°C for 3 h. Samples were then centrifuged at 20,000 × *g* at 4°C for 30 min, and supernatant was removed by pipetting. Beads were rinsed with 500 μL methanol, dried down in a SpeedVac, and resuspended in 95 μL 50 mM (pH 8.4) Tris-formic acid (FA). A total volume of 5 μL trypsin (Sigma Aldrich), reconstituted in 50 mM (pH 8.4) Tris-FA to a final concentration of 0.25 μg/μL, was added for a 6-h tryptic digestion at 37°C with constant vortexing in a thermomixer. Digestion was terminated by adding 1 μL FA, and the supernatant was transferred to a new Eppendorf tube. Beads were then rinsed with 200 μL 0.1% FA, and the supernatant was combined with the previous tube and dried down in a SpeedVac. Digested peptides were resuspended in 50 μL 0.1% FA in 2% acetonitrile (ACN), centrifuged at 20,000 × *g* at 4°C for 30 min, and carefully transferred to LC vials for analysis.

### LC-MS analysis

The LC-MS system consists of a Dionex Ultimate 3000 nano-LC system, a Dinex Ultimate 3000 micro-LC system with a WPS-3000 autosampler, and an Orbitrap Fusion Lumos mass spectrometer. A large-inner diameter (i.d.) trapping column (300-μm i.d. by 5 mm) was implemented before the separation column (75-μm i.d. by 65 cm, packed with 2.5-μm Xselect CSH C_18_ material) for high-capacity sample loading, cleanup, and delivery. For each sample, 8 μL derived peptides was injected for LC-MS analysis. Mobile phases A and B were 0.1% FA in 2% ACN and 0.1% FA in 88% ACN. The 180-min LC gradient profile was 4% for 3 min, 4–11% for 5 min, 11–32% B for 117 min, 32–50% B for 10 min, 50–97% B for 5 min, and 97% B for 7 min and then equilibrated to 4% for 27 min. The mass spectrometer was operated under data-dependent acquisition mode with a maximal duty cycle of 3 s. MS1 spectra was acquired by Orbitrap under 120-k resolution for ions within the *m/z* range of 400–1,500. Automatic gain control and maximal injection time were set at 120% and 50 ms, respectively, and the dynamic exclusion was set at 45 s and ±10 ppm. Precursor ions were isolated by quadrupole using a *m/z* window of 1.2 Th and were fragmented by high-energy collision dissociation for back-to-back Orbitrap/ion trap MS2 acquisition. Orbitrap MS2 spectra were acquired under 15-k resolution with a maximal injection time of 50 ms, and ion trap MS spectra was acquired under rapid scan rate with a maximal injection time of 35 ms. Detailed LC-MS settings and relevant information can be found in a previous publication by Shen et al (69).

### Data processing

LC-MS files were searched against *Cryptococcus neoformans* var. *grubii* serotype A in the Swiss-Prot+TrEMBL protein sequence database (7,429 entries) using Sequest HT embedded in Proteome Discoverer 1.4 (Thermo Fisher Scientific). A target-decoy approach using a concatenated forward and reverse protein sequence database was applied for false-discovery rate (FDR) estimation and control. Searching parameters were as follows: precursor ion mass tolerance, 20 ppm; product ion mass tolerance, 0.8 Da; maximal missed cleavages per peptide, 2; fixed modifications, cysteine carbamidomethylation; and dynamic modifications, methionine oxidation and peptide N-terminal acetylation. Search result merging, protein inference/grouping, and FDR control were performed in Scaffold 5 (Proteome Software, Inc.). For identification, the global protein/peptide FDR was set to 1.0% and at least 2 unique peptides were required for each protein. For quantification, protein abundance was determined by total spectrum counts and total MS2 ion intensities. Results were exported and manually curated in Microsoft Excel.

### Immunoblotting

Cell pellets were lysed in RIPA buffer (supplemented with Halt phosphatase and protease inhibitor cocktail) by bead beating using glass beads. Lysate from the beads was clarified by centrifugation at 20,000 × *g* for 10 min and quantified using Pierce 660-nm protein assay kit (Thermo Fisher Scientific). Ten micrograms of protein was run per sample on 4–15% Mini-Protean TGX stain-free precast gels (Bio-Rad) at 150 V. Proteins were transferred from the gels to nitrocellulose membrane using a Bio-Rad Trans-Blot Turbo transfer system and blocked using Bio-Rad EveryBlot blocking solution for 1 h. Blots were then incubated overnight using 1:1,000 diluted anti-eIF2α (custom made) and anti-p-eIF2α (Abcam, ab3215) antibodies in EveryBlot blocking solution. The signal was detected using horseradish peroxidase-conjugated secondary antibodies with Bio-Rad Clarity Max ECL substrate and imaged on a Bio-Rad ChemiDoc imaging system. Bands were quantified using Image Lab, and normalized total protein was quantified from stain-free gels.

### RNA sequencing

Wild-type and *rmt5*Δ cells were grown to mid-log stage at 37°C and pelleted. Cells were lysed by bead beating, and RNA was extracted using an RNeasy mini kit (Qiagen) and treated using an on-column RNase-free DNase kit (Qiagen). RNA samples were submitted to GeneWiz for library preparation an Illumina sequencing. Two biological replicates were analyzed per strain.

RNA sequencing libraries were prepared using the NEBNext Ultra II RNA library prep kit for Illumina according to the manufacturer’s instructions (NEB). Prior to library prep, mRNAs were enriched with oligo(dT) beads. The sequencing libraries were validated on the Agilent TapeStation (Agilent Technologies) and quantified by using a Qubit 2.0 fluorometer (Invitrogen) as well as by quantitative PCR (KAPA Biosystems). The sequencing libraries were clustered on one lane of a flow cell. After clustering, the flow cell was loaded on the Illumina HiSeq instrument (4000 or equivalent) according to the manufacturer’s instructions. The samples were sequenced using a 2 × 150-bp paired-end configuration.

RNA sequencing reads were trimmed to remove adapter sequences and filtered using Cutadapt. Filtered reads were aligned to the *C. neoformans* H99 genome downloaded from FungiDB using STAR alignment. After mapping, there were approximately 12–14 million reads per sample. The read counts for each gene were calculated using RSEM. Differentially expressed genes between the mutant and wild-type samples were determined using DeSeq2 in R. Following DeSeq2, data were filtered with a 1.75-fold change and an adjusted *p* value threshold of 0.05. Data were visualized using EnhancedVolcano and chromoMap R packages. All RNA sequencing files for this project are available in the NCBI Gene Expression Omnibus (GEO), accession number pending.

## Acknowledgements

This work was funded, in part, by American Heart Association (AHA) grant 33440094 to JCP and AHA predoctoral fellowship 827110 to MCK). We acknowledge Dr. Graham Solomon and Hussain Odeh for assistance in molecular cloning. We acknowledge Dr. Jason Kay (University at Buffalo, Oral Biology) for gifting us the J774A cell line. We thank Dr. Amanda L. M. Bloom for helpful discussions and assistance with the mouse study. We thank Dr. Chelsie Armbruster for providing equipment for spiral plating and colony counting. We thank Dr. Amy Jacobs for providing tissue culture space for macrophage work. We also thank Dr. Karen Dietz for critical review of the manuscript.

## Figure Legends

**Figure S1.**
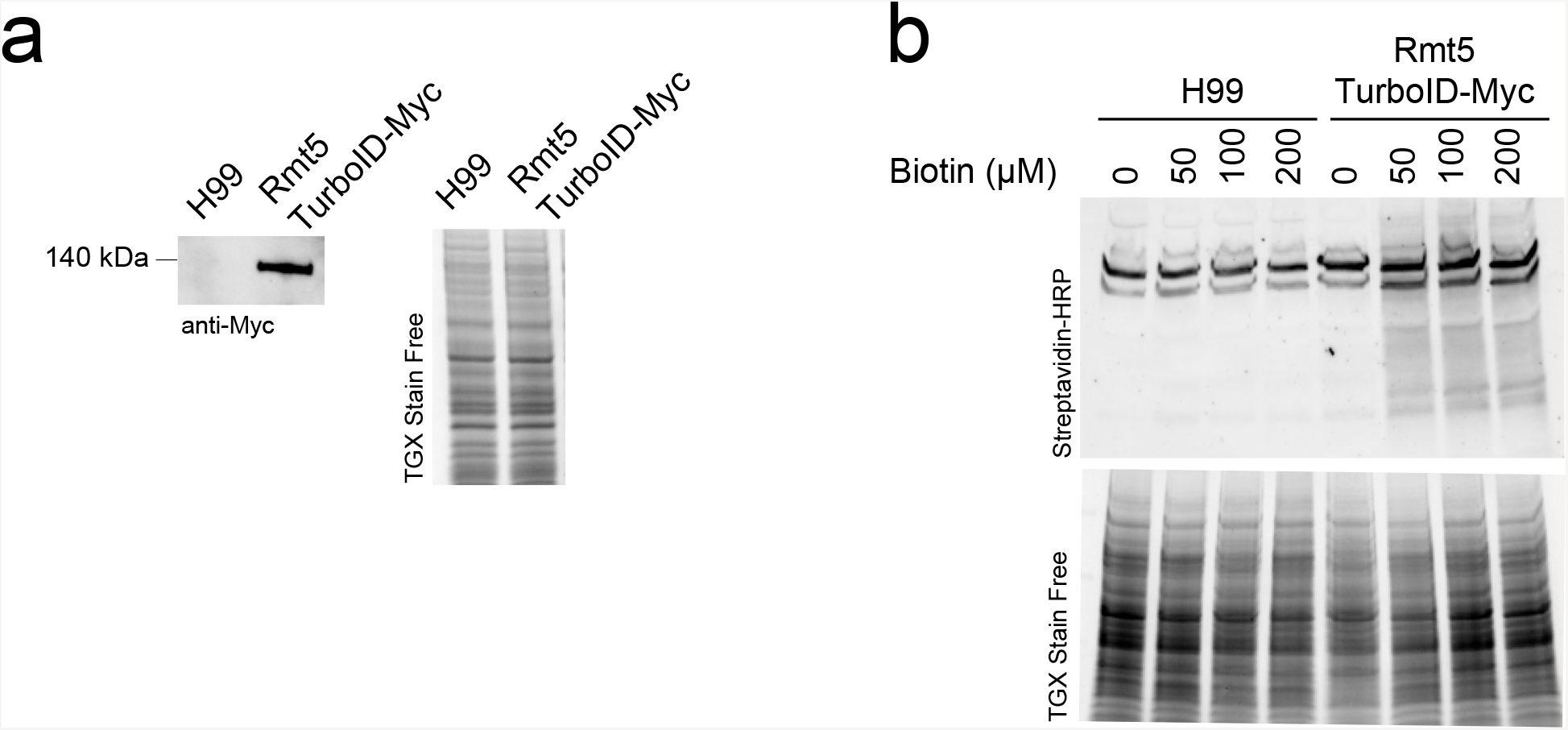
Expression of *RMT5-*TurboID-3×Myc. (**a**) Anti-Myc western blot showing the expression of *RMT5-*TurboID-3×Myc. Stain-free gel is shown as a loading control. (**b**) Titration of biotin and determination of the optimal biotin concentration. Cells were grown in media containing different concentrations of biotin. Western blots probed using streptavidin-HRP show that the 50 µM is optimal to detect biotinylated proteins.

**Figure S2.**
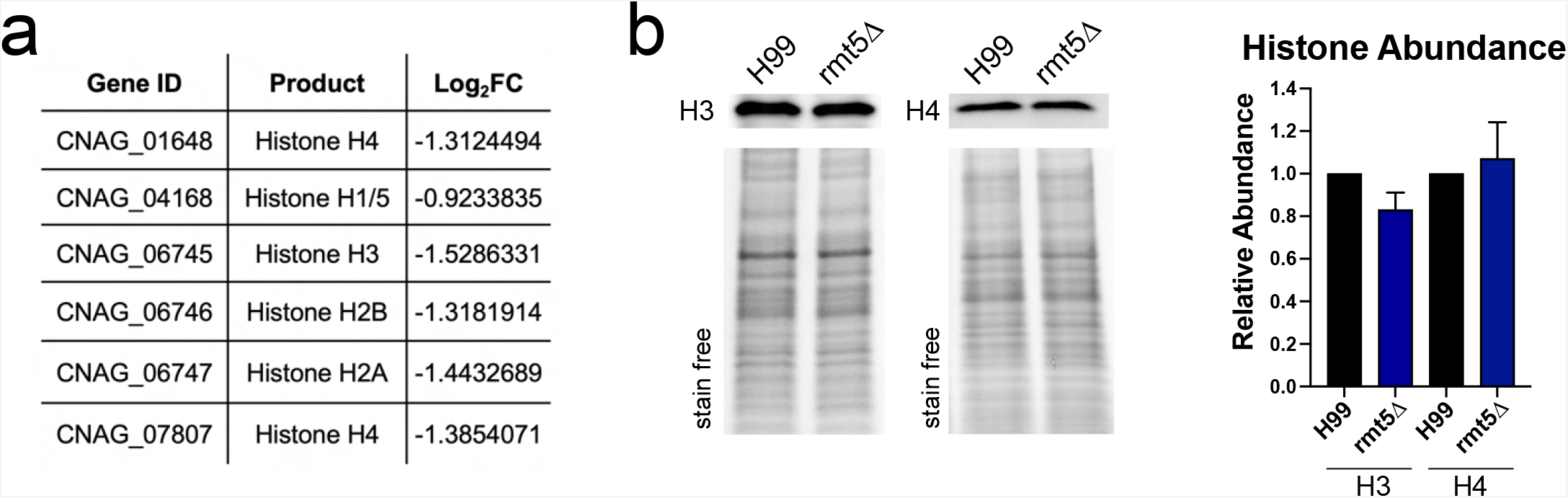
Histone transcripts are downregulated in the *rmt5*Δ mutant. (**a**) RNA-seq log2 fold change of histone transcripts show that they are downregulated in the mutant. (**b**) Western blots showing that the expression of H3 and H4 proteins are not altered in the mutant.

**Figure S3.**
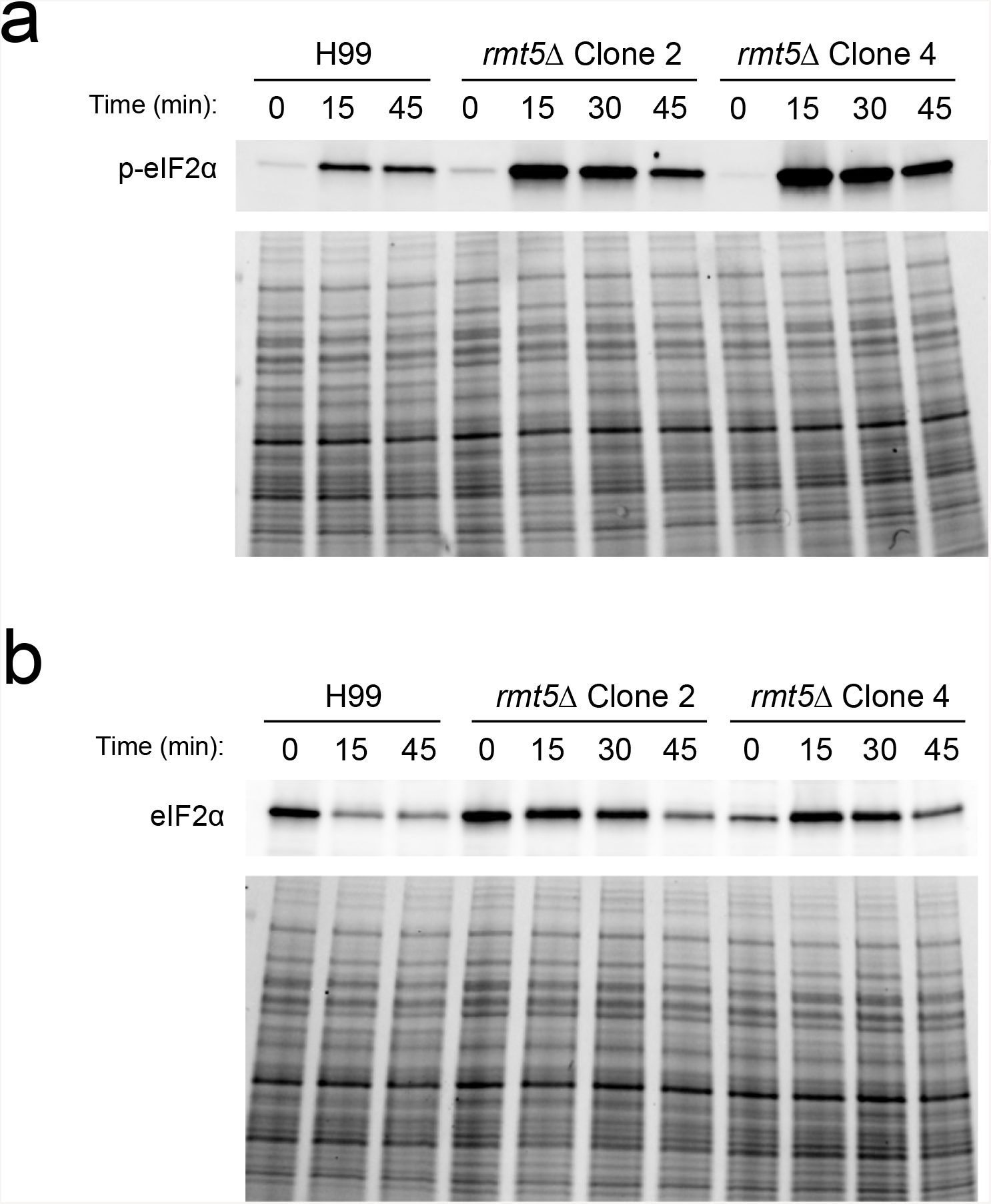
eIF2α abundance and phosphorylation during 3-AT-induced ribosome stalling. New rmt5Δ clones were analyzed for eIF2α phosphorylation (a) and expression (b) using western blotting. Representative images are shown to demonstrate that the clones 2 and 4 have the same eIF2α response to 3-AT as the original mutant.

**Table S1**. Primers used in molecular cloning and cell line construction.

**Table S2**. TurboID protein interaction network.

**Table S3**. TurboID interaction network gene ontology analysis.

**Table S4**. RNA-seq differentially expressed genes.

**Table S5**. RNA-seq gene ontology analysis.

## Notes

### Competing Interest Statement

The authors have declared no competing interest.

## References

1. Conradi C, Shiu A. 2018. Dynamics of Posttranslational Modification Systems: Recent Progress and Future Directions. Biophys J 114:507–515.

2. Dowd CJ, Cooney CL, Nugent MA. 1999. Heparan sulfate mediates bFGF transport through basement membrane by diffusion with rapid reversible binding. J Biol Chem 274:5236–5244.

3. Deribe YL, Pawson T, Dikic I. 2010. Post-translational modifications in signal integration. Nat Struct Mol Biol 17:666–672.

4. Ryšlavá H, Doubnerová V, Kavan D, Vaněk O. 2013. Effect of posttranslational modifications on enzyme function and assembly. J Proteomics 92:80–109.

5. Velázquez-Cruz A, Baños-Jaime B, Díaz-Quintana A, De la Rosa MA, Díaz-Moreno I. 2021. Post-translational Control of RNA-Binding Proteins and Disease-Related Dysregulation. Front Mol Biosci 8:1–15.

6. Wu Q, Schapira M, Arrowsmith CH, Barsyte-Lovejoy D. 2021. Protein arginine methylation: from enigmatic functions to therapeutic targeting. Nat Rev Drug Discov 20:509–530.

7. Maniaci M, Boffo FL, Massignani E, Bonaldi T. 2021. Systematic Analysis of the Impact of R-Methylation on RBPs-RNA Interactions: A Proteomic Approach. Front Mol Biosci 8:1– 19.

8. Liu Q, Dreyfuss G. 1995. In vivo and in vitro arginine methylation of RNA-binding proteins. Mol Cell Biol 15:2800–2808.

9. Yu MC, Mcbride AE, Komili S, Casolari JM, Silver PA. 2004. Arginine methyltransferase affects interactions and recruitment of mRNA processing and export factors 2024–2035.

10. Goulah CC, Read LK. 2007. Differential effects of arginine methylation on RBP16 mRNA binding, guide RNA (gRNA) binding, and gRNA-containing ribonucleoprotein complex (gRNP) formation. J Biol Chem 282:7181–7190.

11. Lott K, Mukhopadhyay S, Li J, Wang J, Yao J, Sun Y, Qu J, Read LK. 2015. Arginine methylation of DRBD18 differentially impacts its opposing effects on the trypanosome transcriptome. Nucleic Acids Res 43:5501–5523.

12. Wei HM, Hu HH, Chang GY, Lee YJ, Li YC, Chang HH, Li C. 2014. Arginine methylation of the cellular nucleic acid binding protein does not affect its subcellular localization but impedes RNA binding. FEBS Lett 588:1542–1548.

13. Guccione E, Richard S. 2019. The regulation, functions and clinical relevance of arginine methylation. Nat Rev Mol Cell Biol 20:642–657.

14. Blanc RS, Richard S. 2017. Arginine Methylation: The Coming of Age. Mol Cell 65:8–24.

15. Hwang JW, Cho Y, Bae GU, Kim SN, Kim YK. 2021. Protein arginine methyltransferases: promising targets for cancer therapy. Exp Mol Med 53:788–808.

16. Lipson RS, Webb KJ, Clarke SG. 2010. Rmt1 catalyzes zinc-finger independent arginine methylation of ribosomal protein Rps2 in Saccharomyces cerevisiae. Biochem Biophys Res Commun 391:1658–1662.

17. Bachand F. 2007. Protein arginine methyltransferases: From unicellular eukaryotes to humans. Eukaryot Cell 6:889–898.

18. Boisvert FM, Côté J, Boulanger MC, Cléroux P, Bachand F, Autexier C, Richard S. 2002. Symmetrical dimethylarginine methylation is required for the localization of SMN in Cajal bodies and pre-mRNA splicing. J Cell Biol 159:957–969.

19. Gao G, Dhar S, Bedford MT. 2017. PRMT5 regulates IRES-dependent translation via methylation of hnRNP A1. Nucleic Acids Res 45:4359–4369.

20. Kang H, Tsygankov D, Lew DJ. 2016. Sensing a bud in the yeast morphogenesis checkpoint: a role for Elm1. Mol Biol Cell 27:1764–1775.

21. Low JKK, Hart-Smith G, Erce MA, Wilkins MR. 2013. Analysis of the proteome of Saccharomyces cerevisiae for methylarginine. J Proteome Res 12:3884–3899.

22. Milliman EJ, Hu Z, Yu MC. 2012. Genomic insights of protein arginine methyltransferase Hmt1 binding reveals novel regulatory functions. BMC Genomics 13:1–14.

23. Yagoub D, Hart-Smith G, Moecking J, Erce MA, Wilkins MR. 2015. Yeast proteins Gar1p, Nop1p, Npl3p, Nsr1p, and Rps2p are natively methylated and are substrates of the arginine methyltransferase Hmt1p. Proteomics 15:3209–3218.

24. Bachand F, Silver PA. 2004. PRMT3 is a ribosomal protein methyltransferase that affects the cellular levels of ribosomal subunits. EMBO J 23:2641–2650.

25. Blum E, Regis A, Conboy A, Clarke S, Elf S, Zurita-Lopez C, McBride AE. 2007. Protein Arginine Methylation in Candida albicans : Role in Nuclear Transport. Eukaryot Cell 6:1119–1129.

26. Li Z, Wu L, Wu H, Zhang X, Mei J, Zhou X, Wang GL, Liu W. 2020. Arginine methylation is required for remodelling pre-mRNA splicing and induction of autophagy in rice blast fungus. New Phytol 225:413–429.

27. Xu X, Chen Y, Li B, Tian S. 2021. Arginine methyltransferase permtc regulates development and pathogenicity of penicillium expansum via mediating key genes in conidiation and secondary metabolism. J Fungi 7.

28. Kalem MC, Subbiah H, Leipheimer J, Glazier VE, Panepinto JC. 2021. Puf4 mediates post-transcriptional regulation of cell wall biosynthesis and caspofungin resistance in cryptococcus neoformans. MBio 12:1–20.

29. Leipheimer J, Bloom ALM, Campomizzi CS, Salei Y, Panepinto JC. 2019. Translational regulation promotes oxidative stress resistance in the human fungal pathogen cryptococcus neoformans. MBio 10:1–13.

30. Bloom ALM, Jin RM, Leipheimer J, Bard JE, Yergeau D, Wohlfert EA, Panepinto JC. 2019. Thermotolerance in the pathogen Cryptococcus neoformans is linked to antigen masking via mRNA decay-dependent reprogramming. Nat Commun 10:1–13.

31. Glazier VE, Kaur JN, Brown NT, Rivera AA, Panepinto JC. 2015. Puf4 regulates both splicing and decay of HXL1 mRNA encoding the unfolded protein response transcription factor in Cryptococcus neoformans. Eukaryot Cell 14:385–395.

32. Kaur JN, Panepinto JC. 2016. Morphotype-specific effector functions of Cryptococcus neoformans PUM1. Sci Rep 6:1–9.

33. Stovall AK, Knowles CM, Kalem MC, Panepinto JC. 2021. A Conserved Gcn2-Gcn4 Axis Links Methionine Utilization and the Oxidative Stress Response in Cryptococcus neoformans. Front Fungal Biol 2:1–14.

34. Lee H, Bien CM, Hughes AL, Espenshade PJ, Kwon-Chung KJ, Chang YC. 2007. Cobalt chloride, a hypoxia-mimicking agent, targets sterol synthesis in the pathogenic fungus Cryptococcus neoformans. Mol Microbiol 65:1018–1033.

35. Chun CD, Liu OW, Madhani HD. 2007. A link between virulence and homeostatic responses to hypoxia during infection by the human fungal pathogen Cryptococcus neoformans. PLoS Pathog 3:0225–0238.

36. Schneider-Poetsch T, Ju J, Eyler DE, Dang Y, Bhat S, Merrick WC, Green R, Shen B, Liu JO. 2010. Inhibition of eukaryotic translation elongation by cycloheximide and lactimidomycin. Nat Chem Biol 6:209–217.

37. Klopotowski T, Wiater A. 1965. Synergism of aminotriazole and phosphate on the inhibition of yeast imidazole glycerol phosphate dehydratase. Arch Biochem Biophys 112:562–566.

38. Guydosh NR, Green R. 2014. Dom34 rescues ribosomes in 3′ untranslated regions. Cell 156:950–962.

39. Hinnebusch AG. 2005. Translational regulation of GCN4 and the general amino acid control of yeast. Annu Rev Microbiol 59:407–450.

40. El Helou G, Hellinger W. 2019. Cryptococcus neoformans Pericarditis in a Lung Transplant Recipient: Case Report, Literature Review and Pearls. Transpl Infect Dis 0–1.

41. Kabir V, Maertens J, Kuypers D. 2018. Fungal infections in solid organ transplantation: An update on diagnosis and treatment. Transplant Rev https://doi.org/10.1016/j.trre.2018.12.001.

42. Malhotra P, Shah SS, Kaplan M, McGowan JP. 2005. Cryptococcal fungemia in a neutropenic patient with AIDS while receiving caspofungin. J Infect 51:181–183.

43. Rajasingham R, Smith RM, Park BJ, Jarvis JN, Govender NP, Chiller TM, Denning DW, Loyse A, Boulware DR. 2017. Global burden of disease of HIV-associated cryptococcal meningitis: an updated analysis. Lancet Infect Dis https://doi.org/10.1016/S1473-3099(17)30243-8.

44. Loyse A, Wilson D, Meintjes G, Jarvis JN, Bicanic T, Bishop L, Rebe K, Williams A, Jaffar S, Bekker LG, Wood R, Harrison TS. 2012. Comparison of the early fungicidal activity of high-dose fluconazole, voriconazole, and flucytosine as second-line drugs given in combination with amphotericin B for the treatment of HIV-associated cryptococcal meningitis. Clin Infect Dis 54:121–128.

45. Chan-Penebre E, Kuplast KG, Majer CR, Boriack-Sjodin PA, Wigle TJ, Johnston LD, Rioux N, Munchhof MJ, Jin L, Jacques SL, West KA, Lingaraj T, Stickland K, Ribich SA, Raimondi A, Scott MP, Waters NJ, Pollock RM, Smith JJ, Barbash O, Pappalardi M, Ho TF, Nurse K, Oza KP, Gallagher KT, Kruger R, Moyer MP, Copeland RA, Chesworth R, Duncan KW. 2015. A selective inhibitor of PRMT5 with in vivo and in vitro potency in MCL models. Nat Chem Biol 11:432–437.

46. Panepinto JC, Komperda KW, Hacham M, Shin S, Liu X, Williamson PR. 2007. Binding of serum mannan binding lectin to a cell integrity-defective Cryptococcus neoformans ccr4Δ mutant. Infect Immun 75:4769–4779.

47. Wei H-H, Fan X-J, Hu Y, Guo M, Fang Z-Y, Wu P, Tian X-X, Gao S-X, Peng C, Yang Y, Wang Z. 2019. A systematic survey of PRMT interactomes reveals the key roles of arginine methylation in the global control of RNA splicing and translation. bioRxiv 746529.

48. Bachand F, Lackner DH, Bähler J, Silver PA. 2006. Autoregulation of Ribosome Biosynthesis by a Translational Response in Fission Yeast. Mol Cell Biol 26:1731–1742.

49. Poornima G, Shah S, Vignesh V, Parker R, Rajyaguru PI. 2016. Arginine methylation promotes translation repression activity of eIF4G-binding protein, Scd6. Nucleic Acids Res 44:9358–9368.

50. Banerjee D, Bloom ALM, Panepinto JC. 2016. Opposing PKA and Hog1 signals control the post-transcriptional response to glucose availability in Cryptococcus neoformans. Mol Microbiol 102:306–320.

51. Musiani D, Bok J, Massignani E, Wu L, Tabaglio T, Ippolito MR, Cuomo A, Ozbek U, Zorgati H, Ghoshdastider U, Robinson RC, Guccione E, Bonaldi T. 2019. Proteomics profiling of arginine methylation defines PRMT5 substrate specificity. Sci Signal 12.

52. Gerstein AC, Fu MS, Mukaremera L, Li Z, Ormerod KL, Fraser JA, Berman J, Nielsen K. 2015. Polyploid titan cells produce haploid and aneuploid progeny to promote stress adaptation. MBio 6:1–14.

53. Semighini CP, Averette AF, Perfect JR, Heitman J. 2011. Deletion of cryptococcus neoformans aif ortholog promotes chromosome aneuploidy and fluconazole-resistance in a metacaspase-independent manner. PLoS Pathog 7.

54. Hu G, Wang J, Choi J, Jung WH, Liu I, Litvintseva AP, Bicanic T, Aurora R, Mitchell TG, Perfect JR, Kronstad JW. 2011. Variation in chromosome copy number influences the virulence of Cryptococcus neoformans and occurs in isolates from AIDS patients. BMC Genomics 12:526.

55. Altamirano S, Fang D, Simmons C, Sridhar S, Wu P, Sanyal K, Kozubowski L. 2017. Fluconazole-Induced Ploidy Change in Cryptococcus neoformans Results from the Uncoupling of Cell Growth and Nuclear Division. mSphere 2:1–18.

56. Chang YC, Lamichhane AK, Kwon-Chung KJ. 2018. Cryptococcus neoformans, unlike candida albicans, forms aneuploid clones directly from uninucleated cells under fluconazole stress. MBio 9:1–14.

57. Sionov E, Lee H, Chang YC, Kwon-Chung KJ. 2010. Cryptococcus neoformans overcomes stress of azole drugs by formation of disomy in specific multiple chromosomes. PLoS Pathog 6:1–13.

58. Stone NRH, Rhodes J, Fisher MC, Mfinanga S, Kivuyo S, Rugemalila J, Segal ES, Needleman L, Molloy SF, Kwon-Chung J, Harrison TS, Hope W, Berman J, Bicanic T. 2019. Dynamic ploidy changes drive fluconazole resistance in human cryptococcal meningitis. J Clin Invest 129:999–1014.

59. Chang YC, Lamichhane AK, Cai H, Walter PJ, Bennett JE, Kwon-Chung KJ. 2021. Moderate levels of 5-fluorocytosine cause the emergence of high frequency resistance in cryptococci. Nat Commun 12:1–13.

60. Basenko E, Pulman J, Shanmugasundram A, Harb O, Crouch K, Starns D, Warrenfeltz S, Aurrecoechea C, Stoeckert C, Kissinger J, Roos D, Hertz-Fowler C. 2018. FungiDB: An Integrated Bioinformatic Resource for Fungi and Oomycetes. J Fungi 4:39.

61. Goppelt A, Steizer G, Lottspeich F, Meisterernst M. 1996. A mechanism for repression of class II gene transcription through specific binding of NC2 to TBP-promoter complexes via heterodimeric histone fold domains 15:3105–3116.

62. Kim I, Sinha S, Maity SN. 1996. Determination of Functional Domains in the C Subunit of the CCAAT-Binding Factor (CBF) Necessary for Formation of a CBF-DNA Complex : CBF-B Interacts Simultaneously with both the CBF-A and CBF-C Subunits To Form a Heterotrimeric CBF Molecule 16:4003–4013.

63. Hartlepp KF, Ferna C, Eberharter A, Gru T, Mu CW, Becker PB. 2005. The Histone Fold Subunits of Drosophila CHRAC Facilitate Nucleosome Sliding through Dynamic DNA Interactions ‡ 25:9886–9896.

64. Abbott J, Marzluff WF, Gall JG. 1999. The Stem-Loop Binding Protein (SLBP1) Is Present in Coiled Bodies of the Xenopus Germinal Vesicle 10:487–499.

65. Gallie DR, Lewis NJ, Marzluff WF. 1996. The histone 3 ′ -terminal stem – loop is necessary for translation in Chinese hamster ovary cells 24:1954–1962.

66. Larochelle M, Bergeron D, Arcand B, Bachand F. 2019. Proximity-dependent biotinylation mediated by TurboID to identify protein-protein interaction networks in yeast. J Cell Sci 132.

67. Wormley FL, Perfect JR. 2005. Immunology of infection caused by Cryptococcus neoformans. Methods Mol Med 118:193–198.

68. Shen S, An B, Wang X, Hilchey SP, Li J, Cao J, Tian Y, Hu C, Jin L, Ng A, Tu C, Qu M, Zand MS, Qu J. 2018. Surfactant Cocktail-Aided Extraction/Precipitation/On-Pellet Digestion Strategy Enables Efficient and Reproducible Sample Preparation for Large-Scale Quantitative Proteomics. Anal Chem 90:10350–10359.

69. Shen X, Shen S, Li J, Hu Q, Nie L, Tu C, Wang X, Orsburn B, Wang J, Qu J. 2017. An IonStar Experimental Strategy for MS1 Ion Current-Based Quantification Using Ultrahigh-Field Orbitrap: Reproducible, In-Depth, and Accurate Protein Measurement in Large Cohorts. J Proteome Res 16:2445–2456.

